# Targeting subtype-specific metabolic preferences in nucleotide biosynthesis inhibits tumor growth in a breast cancer model

**DOI:** 10.1101/2020.04.19.049577

**Authors:** Martin P. Ogrodzinski, Shao Thing Teoh, Sophia Y. Lunt

## Abstract

Investigating metabolic rewiring in cancer can lead to the discovery of new treatment strategies for breast cancer subtypes that currently lack targeted therapies. Using MMTV-Myc driven tumors to model breast cancer heterogeneity, we investigated metabolic differences between two histological subtypes, the epithelial-mesenchymal transition (EMT) and the papillary subtypes, using a combination of genomic and metabolomic techniques. We identified differences in nucleotide metabolism between EMT and papillary subtypes: EMT tumors preferentially use the nucleotide salvage pathway, while papillary tumors prefer *de novo* nucleotide biosynthesis. Using CRISPR/Cas9 gene editing and mass spectrometry-based methods, we determined that targeting the preferred pathway in each subtype resulted in greater metabolic impact than targeting the non-preferred pathway. We further show that knocking out the preferred nucleotide pathway in each subtype has a deleterious effect on *in vivo* tumor growth. In contrast, knocking out the non-preferred pathway has a lesser effect or results in increased tumor growth.

## Introduction

Breast cancer remains the leading cause of cancer-related mortality among women worldwide despite recent trends in decreasing mortality in high income countries^1^, which can be attributed to advances in early detection and treatment^2^. Current treatment strategies for advanced breast cancer often include general chemotherapy and radiation, with the use of targeted therapies, such as endocrine therapy, for specific breast cancer subtypes^3^. These subtypes are often defined based on expression of specific receptors including the estrogen receptor (ER), progesterone receptor (PR), and human epidermal growth factor receptor 2 (HER2), with an additional triple negative breast cancer (TNBC) subtype characterized by the absence of these markers. Breast cancer subtypes can also be classified according to gene expression patterns^4,5^ which often overlap with definitions based on receptor status and other clinical findings^3,5^ and are further able to provide valuable prognostic information^6^. However, targeted therapies are not available for all subtypes of breast cancer, and current rates of recurrence and development of resistance remain problematic^7,8^. It is becoming increasingly clear that breast cancer subtypes have differences in metabolism, and targeting these metabolic pathways could provide new targeted therapy options^9,10^.

Metabolic rewiring is a hallmark of cancer^11^, and significant efforts have been made to identify metabolic vulnerabilities in cancer and leverage these findings to develop novel treatment strategies. Early work defining this concept was performed in the 1920s by Otto Warburg, who observed that tumor cells generally upregulate glycolysis even in aerobic conditions^12^ – a phenomenon now known as the Warburg effect. One of the modern consequences of the Warburg effect is that targeting aerobic glycolysis, by pharmacological inhibition of glycolytic enzymes and by limiting glucose availability through dietary restriction^13-15^, is under investigation as a therapeutic strategy for many types of cancer. However, one of the challenges in using metabolic rewiring to treat cancer arises from the fact that cancer is a remarkably heterogenous disease, and few metabolic vulnerabilities are common to all cancers. This variability is clearly illustrated by breast cancer, which demonstrates heterogeneity on histologic, genetic, and metabolic levels^4,9,16,17^.

In addition to glycolysis, another metabolic pathway commonly targeted in cancer therapy is nucleotide biosynthesis. Nucleotides enable cellular proliferation by facilitating RNA and DNA production^18,19^, and is also required to balance basal rates of RNA turnover in all cells^20^. Nucleotide biosynthesis occurs through two parallel metabolic pathways: 1) *de novo* nucleotide biosynthesis, which generates new nucleotides from precursors derived predominately from glucose and glutamine metabolism and is an energetically costly process, and 2) nucleotide salvage, which allows free bases derived from catabolic processes to be recycled back into nucleotides and is significantly more energetically efficient^20^.

In our current work, we investigate subtype-specific differences in nucleotide metabolism using two histological mouse mammary tumor subtypes derived from the MMTV-Myc mouse tumor model: 1) MMTV-Myc epithelial-mesenchymal-transition (EMT); and 2) MMTV-Myc papillary. This model system mimics the heterogeneity of human breast cancer^21^, and subtypes of the MMTV-Myc model can be correlated with human cancer subtypes based on gene expression patterns: the EMT subtype strongly correlates with the claudin-low subtype, and the papillary subtype correlates more moderately with several human subtypes including basal and luminal breast cancer^22,23^. Since the claudin-low and basal subtypes both have poor prognosis^24,25^, we decided to focus on the corresponding MMTV-Myc EMT and papillary subtypes in this study.

We have previously used cell lines derived from this model system to identify metabolic differences between subtypes^26^. Here, we build on this work by integrating genomic and metabolomic techniques to refine our understanding of the metabolic differences between the EMT and papillary subtypes. We find striking differences in nucleotide metabolism between the two subtypes: the EMT subtype prefers nucleotide salvage pathways, while the papillary subtype prefers *de novo* nucleotide biosynthesis. We further investigate the clinical significance of expressing genes related to *de novo* purine biosynthesis and salvage pathways, and evaluate the consequences of targeting these genes in each subtype using CRISPR/Cas9 gene editing techniques^27,28^. We find that targeting the preferred metabolic pathway of each subtype generally caused the most substantial disruption on nucleotide metabolism and had subtype-specific effects on *in vivo* tumor growth. Notably, targeting the preferred pathway significantly reduced tumor growth while targeting the non-preferred pathway either had no effect on tumor growth, or in some cases significantly increased tumor growth. These results highlight the metabolic heterogeneity of breast cancer subtypes and demonstrate the potential efficacy of tailoring therapies to inhibit subtype-specific metabolism.

## Results

### Metabolite pool sizes and gene expression patterns of MMTV-Myc mouse mammary tumors implicate differences in nucleotide metabolic pathway activity between subtypes

To identify differences in metabolic pathway activities between EMT and papillary mouse mammary tumor subtypes, we integrated a metabolomics analysis with publicly available gene expression data^29^. Metabolites were extracted from flash frozen tumor sections of known histological subtype (Figure 1A) and quantitated using liquid chromatography tandem mass spectrometry (LC-MS/MS). We found metabolites involved in the pentose phosphate pathway (PPP) and metabolites related to nucleotide metabolism to be significantly different between EMT and papillary tumors (Figure 1B; **Supplementary Table 1**). Notably, PPP intermediates including gluconolactone, ribose 5-phosphate, ribulose 5-phosphate, and sedoheptulose phosphate are uniformly elevated in the EMT subtype compared to papillary. The PPP serves several important functions including: 1) production of ribose 5-phosphate which can be used for nucleotide biosynthesis or converted to glycolytic intermediates; 2) production of reducing equivalents in the form of NADPH; and 3) generation of erythrose 4-phosphate which can also be converted to glycolytic intermediates^30^. Several metabolites related to nucleotide metabolism are also different between the EMT and papillary tumors (Figure 1B; **Supplementary Table 1**). For example, EMT tumors have higher levels of inosine monophosphate (IMP), adenine, and inosine compared to papillary tumors. Adenine and inosine are both intermediates in breakdown and salvage pathways of nucleotide metabolism, and IMP is an intermediate for purine biosynthesis. To investigate how these metabolite levels reflect differences in gene expression, we downloaded gene expression data for the EMT and papillary tumors from Gene Expression Omnibus (GEO)^31^ and applied gene set enrichment analysis (GSEA)^32^ using metabolism-related gene sets from the Reactome database^33^. This analysis revealed that genes involved in the PPP (Figure 1C) and nucleobase biosynthesis (Figure 1C) are both significantly enriched for *lower* expression in the EMT subtype compared to the papillary subtype. Therefore, the higher levels of PPP metabolites and IMP observed in EMT tumors (Figure 1B) likely reflect accumulation due to decreased flux through the PPP and nucleobase biosynthesis pathways. Together, this data agrees with our previous *in vitro* findings, where we observed lower nucleotide biosynthesis in EMT cells compared to papillary cells^26^. When considered with our previous results, these findings further demonstrate that EMT and papillary tumors exhibit significant differences in nucleotide metabolism *in vivo*.

**Figure 1.**
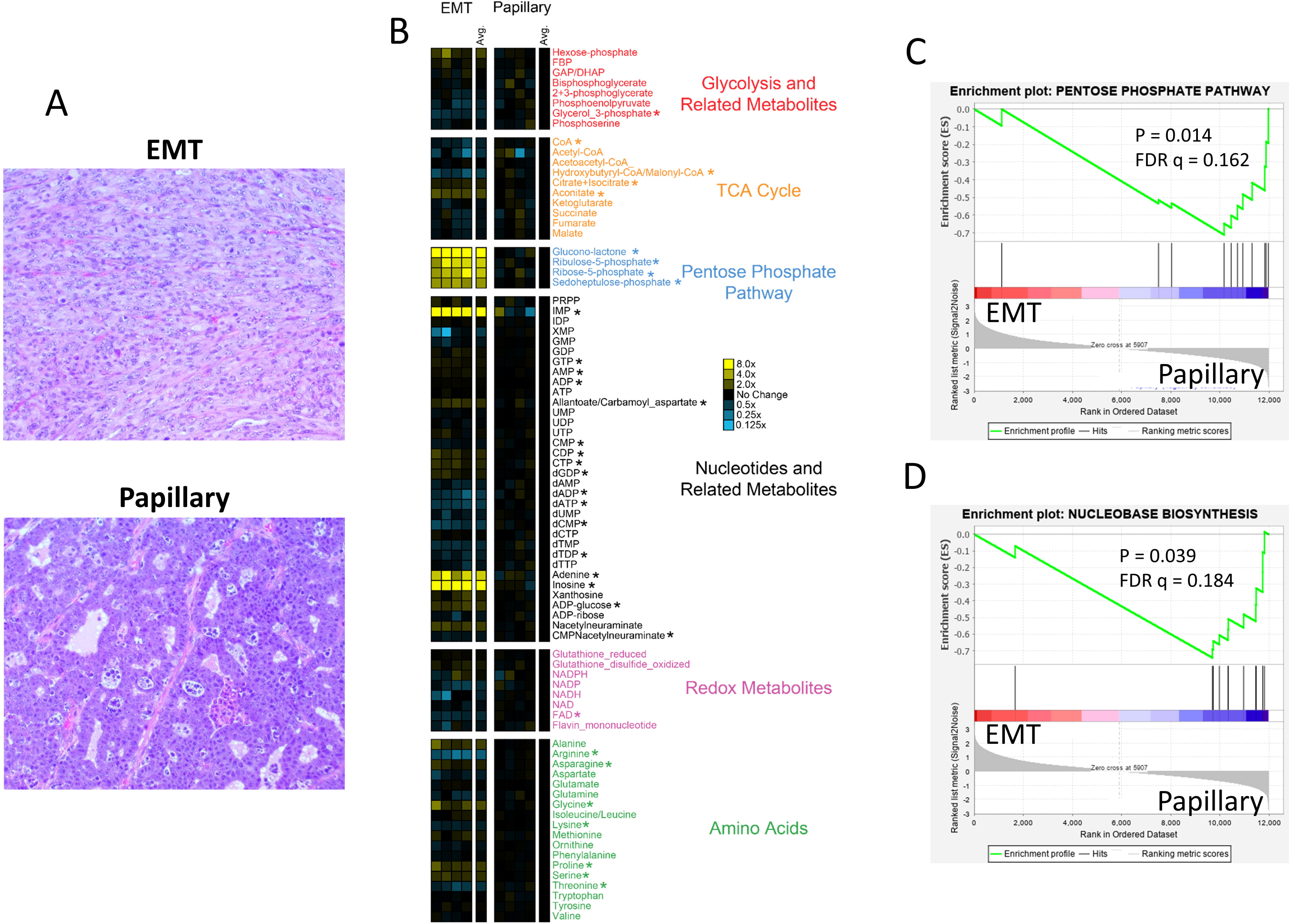
Metabolic profiles and gene expression patterns indicate differences in nucleotide metabolism between MMTV-Myc EMT and papillary tumor subtypes. (**A**) Representative histology images of the EMT and papillary tumor subtypes. (**B**) Heatmap indicating relative metabolite differences between EMT and papillary tumors. Yellow and blue boxes indicate increased or decreased metabolite levels relative to the average of the papillary subtype, respectively. Metabolites with statistically significant differences (p-value < 0.05) are bolded and marked with asterisks (*) Statistical comparisons are listed in **Supplementary Table 1**. (**C**) Gene set enrichment analysis for pentose phosphate pathway genes are significantly enriched (p-value = 0.014, FDR q-value = 0.16) for low expression in EMT tumors vs. papillary tumors. (**D**) Gene set enrichment analysis for genes involved in nucleobase biosynthesis are significantly enriched (p-value = 0.039, FDR q-value = 0.18) for low expression in EMT tumors vs. papillary tumors.

### Expression of nucleotide salvage genes are increased in the EMT subtype

To further characterize gene expression differences in nucleotide metabolism between the EMT and papillary tumors, we used the transcriptome analysis console (TAC) software. We filtered the gene list to include nucleotide metabolism and the PPP genes as denoted within the Reactome database. Based on our GSEA results (Figure 1C-D), we expected genes involved in *de novo* nucleotide biosynthesis and PPP to have higher expression in the papillary subtype. For the EMT subtype, we find that the gene with highest relative expression is *UPP1*, with 18-fold higher expression the EMT subtype vs. the papillary subtype (**Supplementary Table 2**). *UPP1* encodes uridine phosphorylase 1, an enzyme involved in pyrimidine salvage. Together with the observation of higher nucleotide salvage pathway intermediates adenine and inosine in EMT (Figure 1B), this suggests the EMT subtype has higher activity of the nucleotide salvage pathway. Therefore, we decided to focus our analysis on genes involved in *de novo* nucleotide biosynthesis and nucleotide salvage pathways. Hierarchical clustering revealed two major groupings of genes as illustrated by the dendrogram in Figure 2A. The first, smaller group included many nucleotide salvage genes with significantly higher expression in EMT tumors and the second, larger group predominately contained *de novo* biosynthesis genes with significantly lower expression in EMT compared to papillary tumors. Therefore, EMT tumors show a relative preference for nucleotide salvage, while papillary tumors prefer *de novo* biosynthesis. We also considered GSEA results for the nucleotide salvage pathway; however, this gene set as a whole was not significantly enriched for the EMT subtype despite several genes within this pathway having significantly higher expression in the EMT subtype (**Supplementary Figure 1; Supplementary Table 2**). A simplified pathway overview summarizing our results highlights the potential metabolic preferences for nucleotide biosynthesis that are specific to each subtype, with the papillary subtype preferring *de novo* biosynthesis and the EMT subtype preferring nucleotide salvage (Figure 2B).

**Figure 2.**
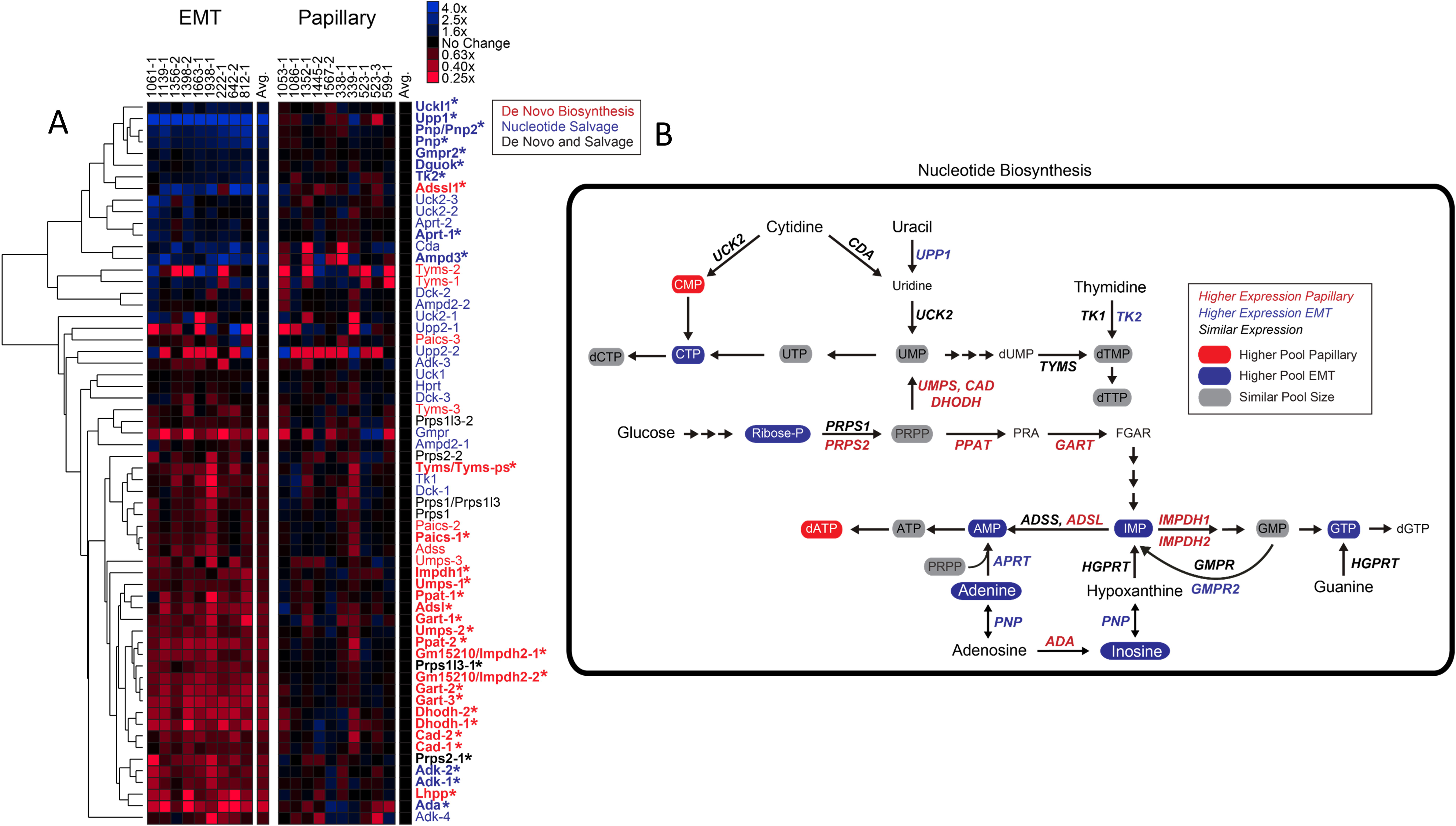
Expression of nucleotide salvage genes is higher in the EMT subtype and expression of *de novo* biosynthesis genes is higher in the papillary subtype. (**A**) Heatmap depicting expression of genes related to nucleotide metabolism. Genes are sorted by hierarchical clustering and color-coded by relationship to nucleotide metabolism pathways. Genes with statistically significant differences (FDR p-value < 0.05) are marked with asterisks (*) Statistical comparisons are listed in **Supplementary Table 2**. (**B**) Summary of nucleotide biosynthesis pathway. Metabolic intermediates and genes are marked according to subtype-specific relationships.

### Expression of key *de novo* and salvage genes are correlated with worse patient outcomes

Our results thus far illustrate the possibility for distinct histological subtypes to utilize different pathways to meet the same metabolic demand for nucleotides. We next sought to determine the potential clinical relevance associated with expression of these genes in human breast cancer. We focused on genes phosphoribosyl pyrophosphate amidotransferase (*PPAT*) and adenine phosphoribosyltransferase (*APRT*) because they encode the rate limiting step for *de novo* purine biosynthesis and salvage of the purine base adenine, respectively, and our findings show that *PPAT* expression is significantly higher in the papillary subtype and *APRT* expression is significantly higher in the EMT subtype (Figure 2B; **Supplementary Table 2**). We used KM plotter, which generates Kaplan-Meier curves using patient data mined from GEO datasets^34^, to generate survival curves with patients stratified by relative gene expression of *PPAT* and *APRT*. We find that in general, patients with high expression of both *PPAT* and *APRT* have worse relapse-free survival (RFS) than patients with low expression of these genes (Figure 3A). This trend is also observed when patients are further divided according to intrinsic subtype. High expression of both *PPAT* and *APRT* is similarly significant for patients with luminal A (Figure 3B) and luminal B (Figure 3C) breast cancer. However, for patients with HER2+ (Figure 3D) and basal (Figure 3E) breast cancer, high expression of *PPAT* is no longer associated with decreased RFS, while high expression of *APRT* remains significant. These results highlight the potential importance of nucleotide metabolism in breast cancer and further suggest that *de novo* purine biosynthesis may be most important for luminal breast cancer, whereas salvage may be relevant for all breast cancer subtypes.

**Figure 3.**
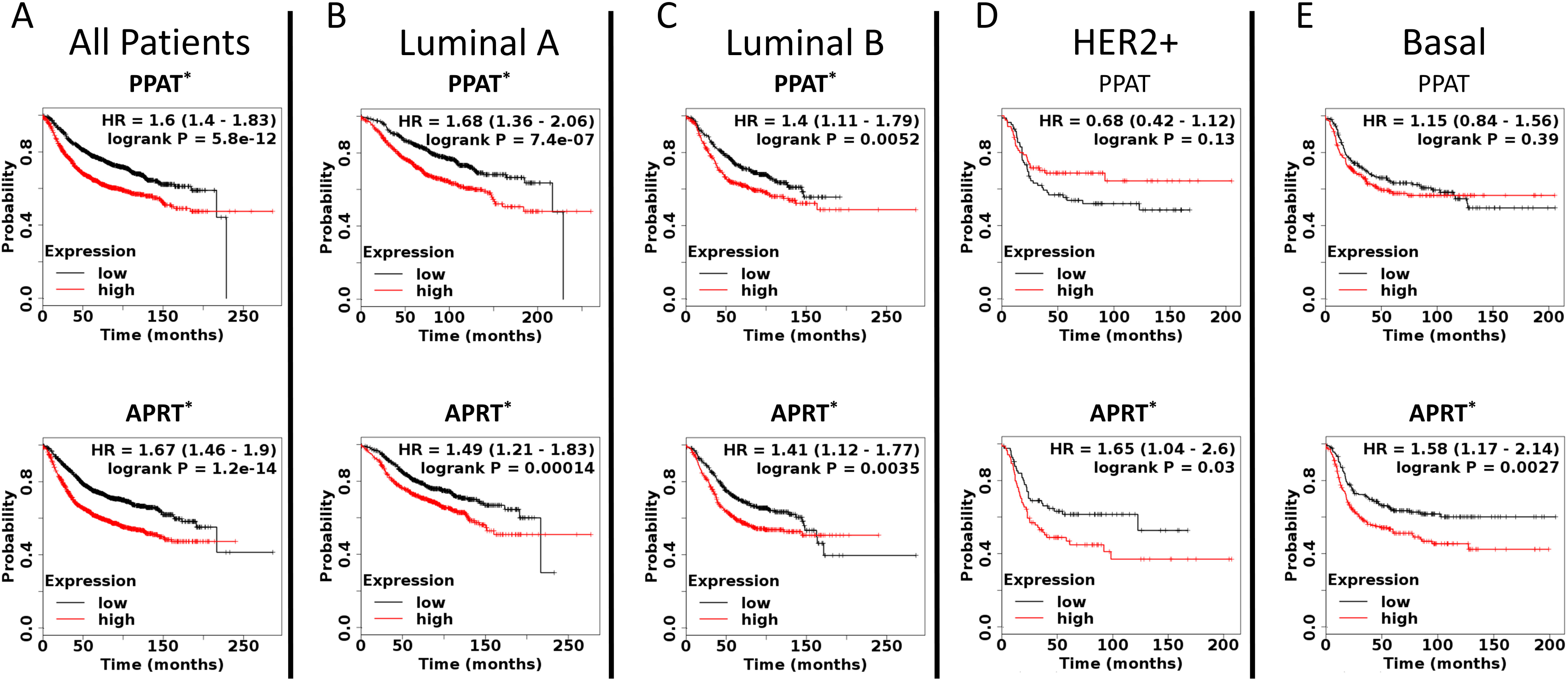
Expression of *de novo* nucleotide biosynthesis gene *PPAT* and nucleotide salvage gene *APRT* are strongly associated with relapse-free survival across breast cancer subtypes. Kaplan-Meier survival curves for (**A**) all breast cancer patients, and (**B-E**) specific breast cancer subtypes. Statistically significant relationships (p-value < 0.05) are bolded and marked with asterisks (*).

### Knocking out *de novo* and salvage genes disrupts cell metabolism in a subtype-specific manner

To further investigate the importance of nucleotide biosynthesis genes *PPAT* and *APRT* in our model, we targeted each gene using CRISPR/Cas9 gene editing^27,28^ in EMT and papillary tumor derived cell lines. We concurrently generated puromycin-resistant control cell lines for each subtype with a non-targeting scramble guide RNA. Clonal lines for each subtype, knockout (KO), and puromycin-resistant scramble control (PSC) were isolated by serial dilution, and successful gene editing was confirmed by Tracking of Indels by Decomposition (TIDE) analysis^35^ (**Supplementary Figure 2**). Western blots were also performed to determine successful KO by protein expression (**Supplementary Figure 3A**). While the APRT antibody worked well, the PPAT bands were inconclusive, with multiple faint bands near the predicted molecular weight of PPAT. We therefore also performed isotope labeling studies to functionally assess how ^13^C-glucose is incorporated into purine biosynthesis in these cell lines. We reasoned that the M-5 isotopologue of ATP represents production from either pathway, since the M-5 isotopologue of ATP is predominately derived from a fully-labeled PRPP molecule with a fully-unlabeled adenine nucleobase, and both *de novo* and salvage pathways utilize PRPP as a substrate. In contrast, the M1-4 and M6-10 isotopologues require labeling of the adenine base, which is only attained through *de novo* biosynthesis. Hence, we distinguish ATP isotopologues as unlabeled (M-0), ATP that may be derived from either *de novo* or salvage pathways (M-5), and ATP that could only be derived from *de novo* biosynthesis (M1-4 and M6-10). Using this approach, we find that, compared to controls, knocking out the salvage gene *APRT* resulted in increased labeling of isotopologues of ATP that can only be derived from *de novo* biosynthesis in both subtypes (Figure 4; **Supplementary Table 3**), which is expected because a larger proportion of ATP is now derived from *de novo* biosynthesis instead of salvage. Additionally, targeting the *de novo* biosynthesis gene *PPAT* caused significantly decreased labeling of ATP isotopologues derived from *de novo* biosynthesis (Figure 4; **Supplementary Table 3**). These results provide strong evidence that the metabolic activity of nucleotide salvage and *de novo* biosynthesis have been significantly decreased in the *APRT* and *PPAT* KO cell lines, respectively.

**Figure 4.**
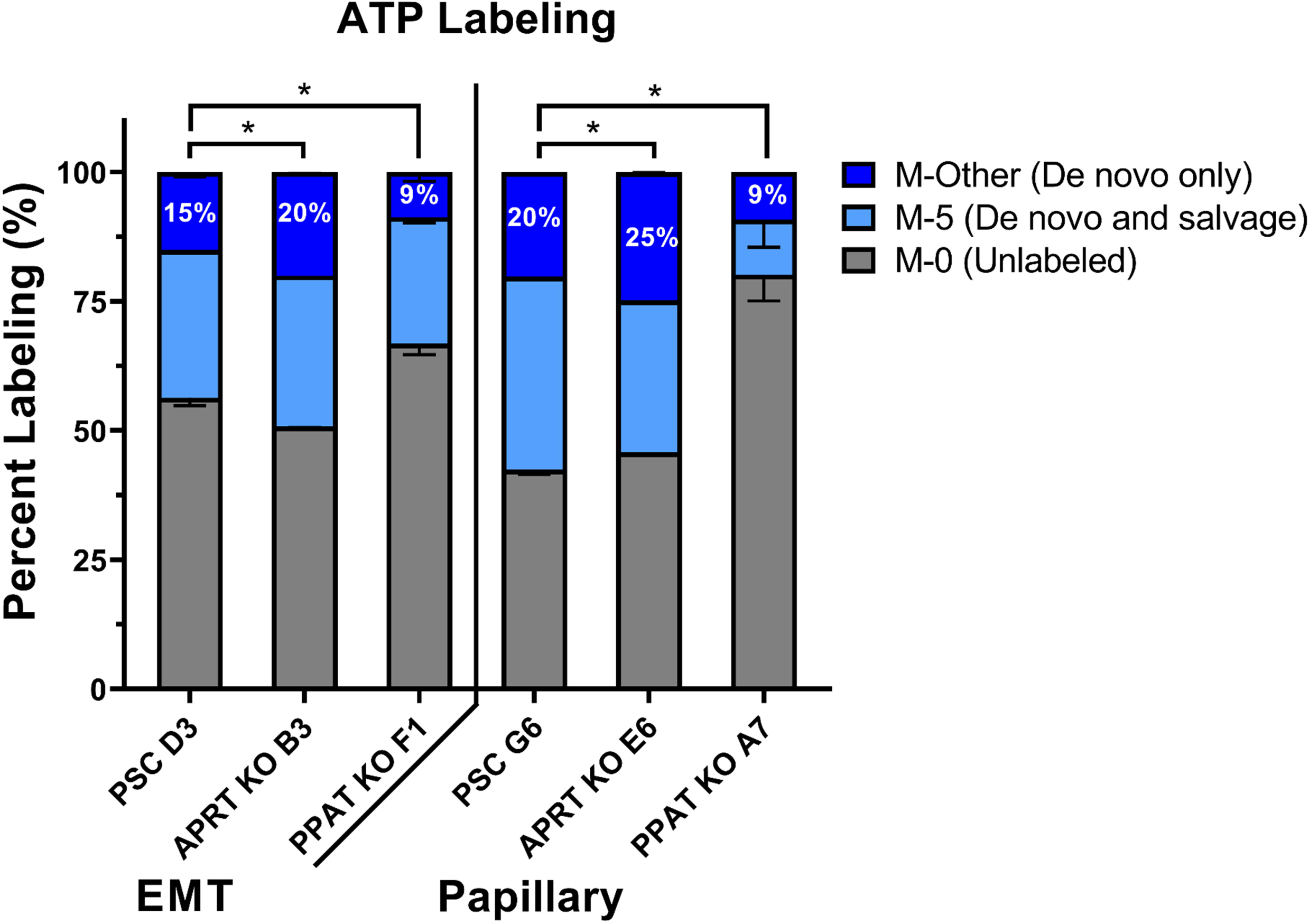
13C-Isotope incorporation from glucose into ATP biosynthesis is altered after targeting *de novo* and salvage genes. Grey boxes represent the unlabeled (M-0 isotopologue) proportion of ATP. Light blue boxes represent the M-5 isotopologue, which can be derived from either *de novo* or salvage pathways. Dark blue boxes represent the sum of all other isotopologues of ATP (M1-4 and M6-10), which are derived from *de novo* ATP biosynthesis. Data are displayed as means ± S.D., N = 3 (*p-value < 0.05). Statistical comparisons are listed in **Supplementary Table 3**.

We also analyzed the abundance of a wide range of metabolites in CRISPR edited cell lines relative to the control line of each respective subtype using the targeted LC-MS/MS method described above. We found significant differences between cell lines across several metabolic pathways (**Supplementary Figure 4; Supplementary Table 4**). As expected, the most consistently altered metabolites include PPP related metabolites (Figure 5A), nucleoside triphosphates (NTPs; Figure 5B), and deoxynucleoside triphosphates (dNTPs; Figure 5C). Notably, the relative abundance of these metabolites is generally most different when the preferred metabolic pathway for each subtype has been targeted. For example, ATP (Figure 5B) and dATP (Figure 5C) levels are significantly decreased in the papillary *PPAT* KO cell line compared to the papillary control and *APRT* KO lines, while the EMT *PPAT* KO is similar to EMT control for these metabolites. Additionally, the papillary *PPAT* KO cells have lower levels of most PPP intermediates, but significantly higher levels of phosphoribosyl pyrophosphate (PRPP; Figure 5A) which is used by both *de novo* biosynthesis and salvage pathways and is produced from the PPP intermediate ribose-5-phosphate. Most NTPs and dNTPs are also decreased in the papillary *PPAT* KO line compared to the control or *APRT* KO line (Figure 5B-C). The exception to this is the pyrimidine uridine triphosphate (UTP), which is significantly increased in the papillary *PPAT* KO compared to control and *APRT* KO lines (Figure 5B). Together, these results show that targeting *PPAT* in the papillary cells creates a metabolic bottleneck by blocking *de novo* purine biosynthesis, causing: 1) decreased levels of intermediates in the PPP, the feeder pathway into nucleotide biosynthesis; and 2) increased levels of PRPP which would normally function to increase *de novo* purine and pyrimidine biosynthesis through feedforward mechanisms^20^. This PRPP-driven feedforward mechanism would also increase *de novo* pyrimidine biosynthesis in the papillary *PPAT* KO cells and explain the increased UTP levels observed in papillary *PPAT* KO cells (Figure 5B).

**Figure 5.**
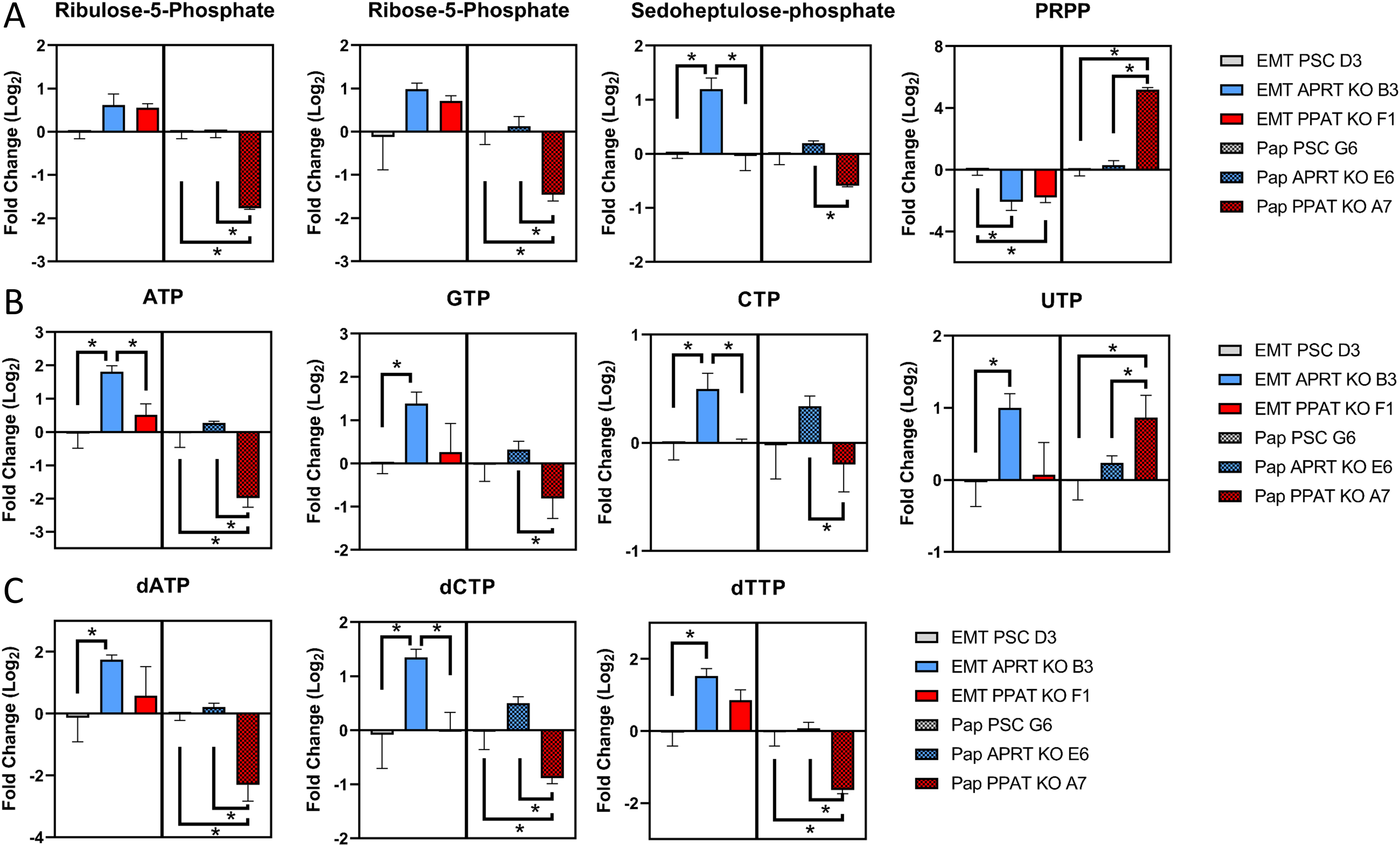
Metabolite levels are most affected by targeting the preferred nucleotide biosynthetic pathway for each subtype. The abundance of (**A**) metabolites related to the pentose phosphate pathway and (**B-C**) nucleotides are most altered within each subtype when *APRT* is knocked out in EMT (left half of each graph) and when *PPAT* is knocked out in papillary (right half of each graph). Data are displayed relative to the control for each subtype and represent means ± S.D., N = 3 (*p-value < 0.05). Statistical comparisons are listed in **Supplementary Table 4**.

Decreased PPP intermediates, increased PRPP, and alterations in NTP and dNTP levels are not observed in the EMT *PPAT* KO cells, likely due to the metabolic preference of the EMT subtype to salvage nucleotides. Indeed, targeting the salvage pathway caused significant metabolic alterations in the EMT *APRT* KO cells compared to the control: higher levels of most nucleotides (Figure 5B-C) suggest that EMT *APRT* KO cells are forced to switch to *de novo* biosynthesis when their preferred means of obtaining nucleotides via salvage is inhibited. In the papillary *APRT* KO line, nucleotide levels are not significantly changed from control levels (Figure 5B-C). Taken together, the above results indicate that the greatest impact on nucleotide metabolism is achieved when the preferred nucleotide biosynthesis pathway of each subtype is inhibited, while inhibiting the non-preferred pathway has minimal effects.

### Targeting nucleotide *de novo* biosynthesis and salvage genes impact tumor growth in a subtype-specific manner

To determine the *in vivo* effects of targeting the preferred nucleotide biosynthesis pathway for each subtype, we monitored tumor growth of KO and control cell lines injected in mice. Control or KO cells were first injected into the mammary fat pad of syngeneic mice to generate tumors, then the resulting tumors were resected, and fragments of these tumors were orthotopically implanted into new cohorts of mice to monitor tumor growth over time. This was performed because implantation of tumor fragments, rather than direct injection of tumor cells, resulted in less variability in the lag time of tumor growth. As expected, the EMT tumors grew slowest when the preferred nucleotide salvage pathway gene *APRT* is targeted: EMT *APRT* KO tumors were significantly smaller (762.8 ± 108.4 mm^2^, n = 5) at 24 days post implantation as compared to the *PPAT* KO tumors (982.7 ± 116.1 mm^2^, n = 5). The EMT *PPAT* KO tumors also grew slower than the PSC tumors (1344.6 ± 141.7 mm^2^, n = 6), which were the largest at 24 days post implantation (Figure 6A; **Supplementary Table 5**). Consistent with the reliance of the papillary subtype on *de novo* nucleotide biosynthesis, targeting *PPAT* prevented papillary cells from growing tumors *in vivo* (Figure 6B; **Supplementary Table 5**). Surprisingly, targeting the non-preferred nucleotide salvage gene *APRT* caused papillary tumors to grow larger (1161.8 ± 155.8 mm^2^, n = 5) than the PSC tumors (514.0 ± 114.0 mm^2^, n = 5) at 24 days post implantation. Taken together, these results indicate that *de novo* nucleotide biosynthesis is a critical metabolic pathway for papillary tumors, and further demonstrate targeting a non-preferred metabolic pathway could have the unintended side effect of increasing tumor growth.

**Figure 6.**
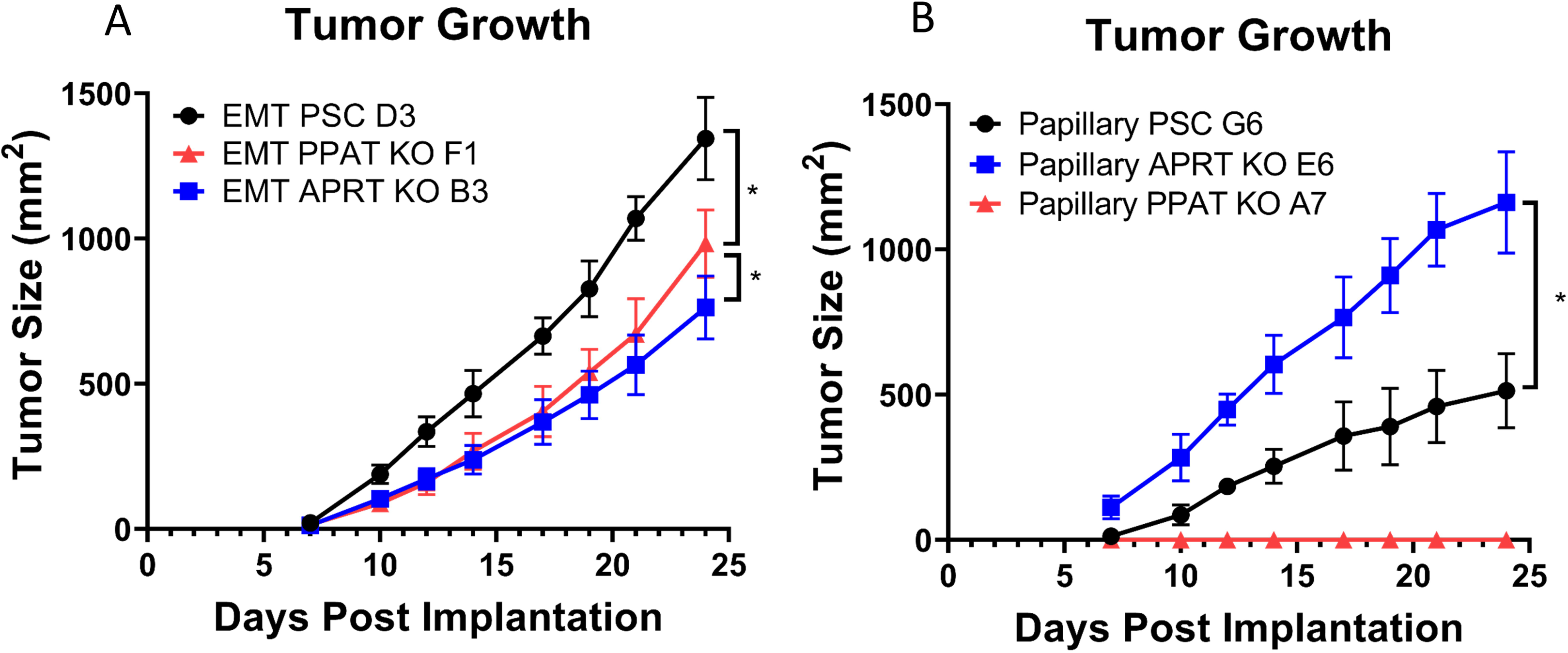
Tumor growth for each subtype is decreased after knocking out the preferred nucleotide metabolism pathway. In vivo growth curves for (**A**) EMT and (**B**) papillary tumors. Data are displayed as means ± S.D. (*p-value < 0.05). Statistical comparisons are listed in **Supplementary Table 5**.

To determine whether differences in tumor sizes are attributable to changes in proliferation or cell death, immunohistochemical (IHC) analysis was performed. Ki67 staining and terminal deoxynucleotidyl transferase dUTP nick end labeling (TUNEL) assays were performed to measure proliferation and necrosis within the tumors, respectively. As expected, Ki67 staining is directly proportional to tumor growth in each subtype. The EMT PSC tumors have significantly more Ki67^+^ nuclei than both KOs, and the EMT *APRT* KO tumors have significantly fewer compared to *PPAT* KOs (Figure 7A; **Supplementary Table 6**). This indicates that the *APRT* KO EMT tumors grow slower due to decreased proliferation. In papillary tumors, the *APRT* KOs have significantly more Ki67^+^ nuclei than the PSC tumors (Figure 7B; **Supplementary Table 6**), showing these tumors grow more quickly due to increased proliferation. TUNEL assays show that in the EMT subtype, both KOs were significantly more necrotic than the control tumors (Figure 7C; **Supplementary Table 7**). In the papillary subtype, no difference in staining was observed between control and *APRT* KO tumors (Figure 7D; **Supplementary Table 7**), indicating that the observed differences in tumor growth are not due to differences in tumor necrosis.

**Figure 7.**
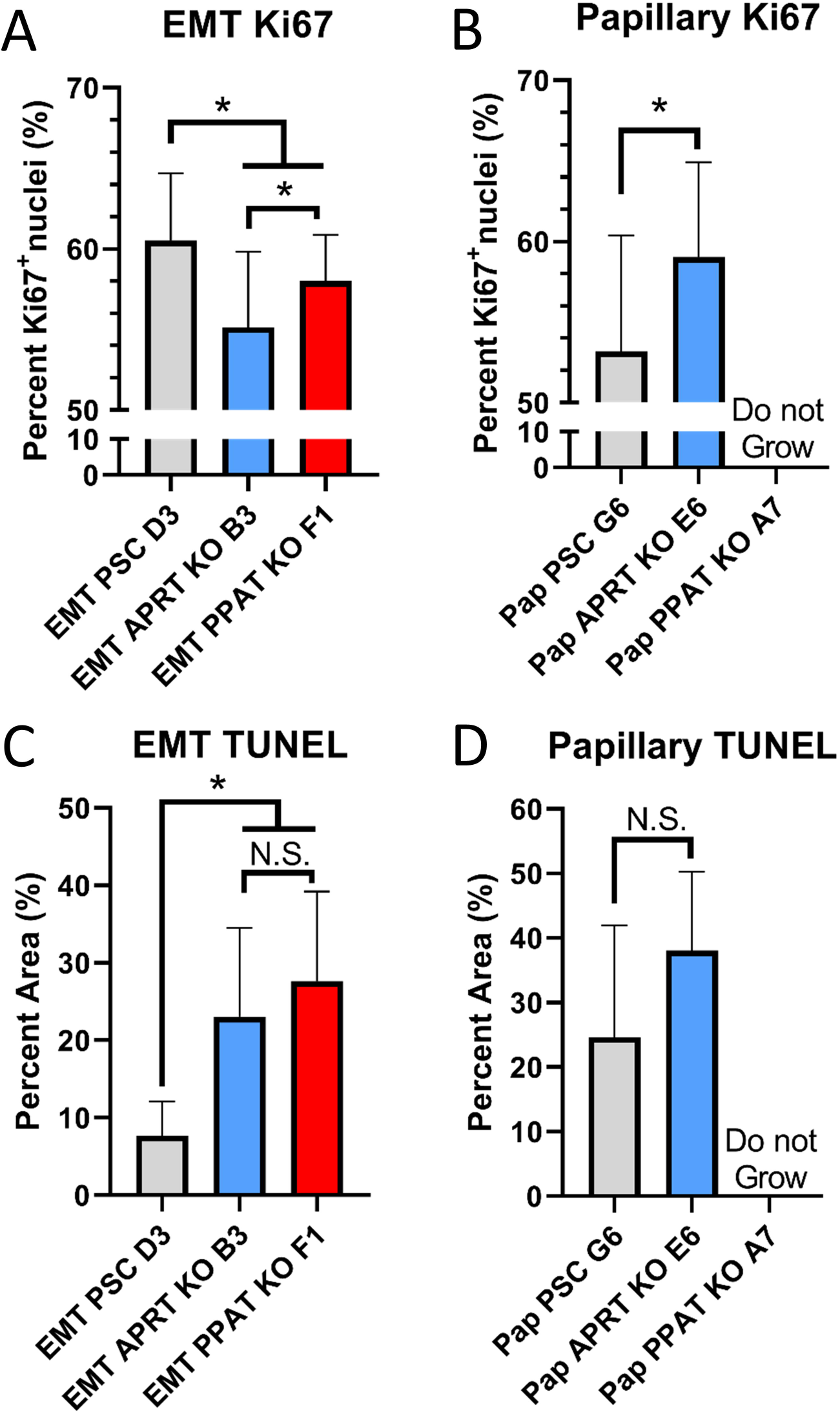
IHC analysis reveals decreased proliferation in slower growing tumors. IHC analysis for Ki67 staining in (**A**) EMT and (**B**) papillary tumors as well as TUNEL assay for (**C**) EMT and (**D**) papillary tumors. Data are displayed as means ± S.D. (*p-value < 0.05). Statistical comparisons are listed in **Supplementary Tables 6-7**.

To validate the monoclonal tumor growth findings, the tumor growth of additional clones for each subtype were measured (**Supplementary Figure 5; Supplementary Table 5**). Four additional papillary *PPAT* KO clones were tested with ATP labeling comparable to Figure 4, as well as one additional EMT *PPAT* KO clone (**Supplementary Figure 6**; **Supplementary Table 8**). Two additional confirmed *APRT* KO clones for each subtype were also injected into mice for *in vivo* testing (**Supplementary Figure 3B**). To validate the controls, the tumor growth of an additional PSC clone and wild-type tumors of each subtype were measured. The clonal PSC tumors for the EMT subtype grew similarly and are larger than the EMT wild-type tumors at 24 days post implantation, indicating the clonal selection process may select for more aggressive clones of this subtype. Additionally, the two EMT *PPAT* KOs grew comparably and were also similar in size to the wild-type EMT tumors. However, the additional two EMT *APRT* KO clonal cell lines failed to generate tumors (**Supplementary Figure 5A; Supplementary Table 5**). For the papillary subtype, one *APRT* KO clone again grew more quickly than the control tumors, while the remaining clone grew similarly to the PSC and wild-type papillary tumors (**Supplementary Figure 5B; Supplementary Table 5**). Consistent with the results shown in Figure 6, the four additional papillary *PPAT* KO clonal cell lines also failed to generate tumors. Taken together, our findings demonstrate the importance of targeting subtype-specific metabolic vulnerabilities to effectively control tumor growth. In addition, inhibiting a non-preferred metabolic pathway not only fails to reduce tumor growth, but can have the detrimental effect of increasing tumor growth.

## Discussion

In this study, we used a combination of genomic and metabolomic techniques to identify subtype-specific metabolic preferences in nucleotide metabolism in the EMT and papillary tumor subtypes derived from the MMTV-Myc mouse model. We discovered that the EMT subtype prefers nucleotide salvage whereas the papillary subtype relies on *de novo* nucleotide biosynthesis. We also investigated patient outcomes and identified that high expression of the nucleotide salvage gene *APRT* is correlated with worse RFS across breast cancer subtypes, while high expression of the *de novo* biosynthesis gene *PPAT* is associated with worse outcomes in patients with luminal breast cancer. We further characterized the metabolic effects of targeting both the preferred and non-preferred pathway in the EMT and papillary subtypes and demonstrate the effect of knocking out these pathways on the *in vivo* tumor growth of each subtype.

Our results demonstrate that targeting the preferred metabolic pathway for nucleotide biosynthesis reduces tumor growth in both EMT and papillary tumors. Sustained proliferation is a hallmark of cancer^11^, and to achieve this cancer cells have a high requirement for nucleotide biosynthesis. Indeed, targeting nucleotide metabolism has long been used as a staple of cancer therapy with early examples including the folate analog methotrexate (MTX) and the pyrimidine analog 5-fluorouracil (5FU). MTX inhibits dihydrofolate reductase and blocks one-carbon metabolism that is essential for several *de novo* biosynthetic reactions^36^, and 5FU inhibits thymidylate synthase, which catalyzes the *de novo* production of thymidine monophosphate^37^. Other compounds targeting *de novo* nucleotide biosynthesis including 6-mercaptopurine^38^, leflunomide^39^ and brequinar^40^ are also currently approved or under investigation as cancer therapeutics. Notably for our model, an active metabolite of 6-mercaptopurine inhibits PPAT^38^ and could prove effective at inhibiting growth of the papillary subtype. However, 6-mercaptopurine and several other *de novo* nucleotide metabolism-targeting compounds including 5FU and gemcitabine are activated by the nucleotide salvage pathway, and downregulation of this pathway could provide a potential resistance mechanism to these compounds^41,42^.

In our model, the papillary subtype has increased MYC signaling compared to the EMT subtype^29^, and its metabolic preference for *de novo* nucleotide biosynthesis highlights the role of MYC as a master regulator of nucleotide biosynthesis^43,44^. MYC amplification is a common feature of many human cancers^45^ and occurs in 15.7% of breast cancers^46^. In TNBC specifically, it has been shown that chemotherapy with doxorubicin adaptively upregulates *de novo* pyrimidine biosynthesis and co-treatment of TNBC xenografts with doxorubicin and the *de novo* pyrimidine biosynthesis inhibitor leflunomide is more effective at treating TNBC tumors than doxorubicin alone^47^. Upregulated *de novo* purine biosynthesis, directed by MYC signaling, has also been implicated as a key metabolic pathway in glioblastoma and targeting *de novo* purine biosynthesis genes improved survival and reduced tumor burden in an *in vivo* model of glioblastoma^48^. These studies and ours strongly suggest that further development of compounds targeting *de novo* nucleotide biosynthesis will be useful to treat many types of cancer.

Our results show that targeting nucleotide salvage also attenuates tumor growth in a subset of cancers that prefer this pathway, such as the EMT subtype MMTV-Myc tumors (Figure 6A). The EMT subtype has previously been correlated with the claudin-low subtype of human breast cancer based on gene expression patterns^22,23^, and additional studies should be performed to determine whether nucleotide salvage is also a metabolic vulnerability in claudin-low breast cancer. Nucleotides and related metabolites are abundant in the extracellular space and serve important biological functions: purines play a significant role as signaling molecules^49^, and pyrimidine release by tumor-associated macrophages has been shown to mediate gemcitabine resistance in animal models of pancreatic cancer^50^. Therefore, the uptake and utilization of these metabolites should be further investigated as therapeutic targets. Unfortunately, there are currently very few available drugs that target nucleotide salvage. Two indirect examples of salvage inhibitors are dilazep and dipyridamole. These compounds act through inhibition of equilibrative nucleoside transporters (ENTs) and function as vasodilators, prevent platelet aggregation, and are currently approved to treat cardiovascular disease^51^. ENT inhibition indirectly blocks nucleotide salvage pathways by preventing uptake of nucleosides and nucleobases. ENTs also mediate the uptake of nucleoside analogs like gemcitabine^52^, which means ENT inhibition as a means to block nucleotide salvage would not be compatible for combination therapy with these drugs. Our findings support the development of therapeutic compounds to specifically target nucleotide salvage pathways. This could prove particularly beneficial for patients diagnosed with claudin-low breast cancer, which carries a poor prognosis and does not have targeted therapies^24^.

Our results further reveal the concerning possibility that targeting a non-preferred pathway can cause an increase in tumor growth. Specifically, when *APRT* was targeted in the papillary subtype, two of three clones grew tumors surprisingly fast, while the remaining clone grew comparably to control tumors (Figure 6 and **Supplemental Figure 9)**. In our current study, we used tumors and cell lines from histologically pure samples; however, this is not always the case in spontaneous tumors. Specifically regarding the MMTV-Myc mouse model, spontaneous tumors develop with a wide variety of histologies, including mixed tumors composed of multiple subtypes in one region^29^. If we consider a possible mixed tumor that is predominately EMT with a minor papillary component, our results indicate that treating it by inhibiting nucleotide salvage alone would likely be ineffective for the papillary component and could even have the unintended side effect of increasing the growth of the papillary portion of the tumor. One implication of this finding is that, for a mixed tumor exhibiting both EMT and papillary histologies, it may be safer to target *de novo* biosynthesis rather than the salvage pathway, because while EMT subtype cells prefer salvage, blocking *de novo* biosynthesis still has a small inhibitory effect on tumor growth (Figure 6A); on the other hand, blocking the salvage pathway in papillary subtype cells can have the opposite and undesirable effect of *increasing* tumor growth (Figure 6B).

In human breast cancer, intratumor heterogeneity can manifest in many ways, including on morphologic and genomic levels^53^. The importance of this heterogeneity is particularly notable when considering biomarker expression; for example, current recommendations report a positive finding if at least 1% of tumors cells are positive for the estrogen receptor (ER)^54^. Since the degree of ER positivity is also directly correlated with patient outcomes following anti-endocrine treatment^55^, it is clear that the intratumor heterogeneity of this biomarker has important clinical implications. Based on our present findings, metabolic vulnerabilities can be used to design new treatments for breast cancer subtypes. However, the possibility of inadvertently stimulating tumor growth by improperly targeting metabolism should also be considered further, especially in recognition of the significant heterogeneity of breast cancer. Further work should be directed at determining whether subtypes of human breast cancer, which are known to exhibit different metabolic features^9^, have differences in metabolic vulnerabilities, and whether targeting non-preferred pathways is detrimental.

In conclusion, our findings demonstrate that distinct histologic subtypes of breast cancer exhibit different metabolic vulnerabilities in terms of their preferred nucleotide biosynthesis pathways, and that inhibiting the preferred pathway greatly impacts metabolism as well as *in vivo* tumor growth. Crucially, we also show that targeting the non-preferred pathway is not only less effective in controlling tumor growth but may have the opposite effect of increasing tumor growth. Our results underscore a critical need to elucidate the distinct metabolic preferences of different breast cancer subtypes in order to design effective targeted therapies for each subtype.

## Methods

### Primary mouse tumors

All animal use was performed in accordance with institutional and federal guidelines. Primary MMTV-Myc EMT and MMTV-Myc papillary tumors were acquired as a gift from Dr. Eran Andrechek and have been previously described^29^. Tumors were sectioned, formalin-fixed, and paraffin embedded for histological examination with hematoxylin and eosin staining. Wild-type EMT and papillary tumors were cryopreserved in a mixture of 90% FBS and 10% DMSO. Tumor derived cell lines were established by mechanical dissociation of primary tumors using scissors, followed by culturing tumor pieces in cell culture media^56^.

### Metabolic profiling

Unlabeled, targeted metabolomics was performed as previously described^57^. Briefly, cells were seeded in 6-well tissue culture plates at 50,000 cells/well and cultured for 48 hours. Cells were washed with saline (VWR, Radnor, Pennsylvania, 16005-092) and metabolism was quenched by addition of cold methanol. Flash frozen tumor tissue was pulverized using a liquid nitrogen cooled mortar and pestle and cold methanol and water was added to the tissue sample. The tissue samples were further processed using a Precellys Evolution homogenizer (Bertin Instruments) operating a single 10s cycle at 10000 rpm. Extracts were then transferred to 1.5 ml Eppendorf tubes and cold chloroform was added to each tube and vortexed for 10 minutes at 4°C. The final metabolite extraction solvent ratios were methanol:water:chloroform (5:2:5). The polar phase was collected and dried under a stream of nitrogen gas. The dried metabolites were then resuspended in HPLC-grade water for analysis. LC-MS/MS analysis was performed with ion-pairing reverse phase chromatography using an Ascentis Express column (C18, 5 cm x 2.1 mm, 2.7 µm, MilliporeSigma, 53822-U) and a Waters Xevo TQ-S triple quadrupole mass spectrometer. Mass spectra were acquired using negative mode electrospray ionization operating in multiple reaction monitoring (MRM) mode. Peak processing was performed using MAVEN^58^ and data for each sample was normalized to the mean signal intensity for all metabolites in the analysis. Metabolites were grouped by relationship to metabolic pathways. Heatmaps were generated using Cluster 3.0^59^ and exported using Java Treeview^60^.

### Gene expression analysis

Gene expression data for MMTV-Myc EMT and papillary data was downloaded from GEO using accession number GSE15904. The following EMT CHP datasets were downloaded: GSM399180, GSM399202, GSM399204, GSM399217, GSM399226, GSM399235, GSM399238, GSM399252, and GSM399259. The following papillary CHP datasets were downloaded: GSM399183, GSM399184, GSM399196, GSM399197, GSM399200, GSM399216, GSM399222, GSM399234, GSM399241, and GSM399245. Gene set enrichment analysis^32^ was performed by converting gene expression data to the required file formats and using the GSEA software available to download from www.gsea-msigdb.org/gsea/index.jsp. Reactome^33^ metabolism gene sets were identified as all participant and sub-participant gene sets under the Reactome Metabolism pathway (stable identifier R-HSA-1430728) and were downloaded from the MSigDB Canonical pathways collection^61^. Differential gene expression was determined using Transcriptome Analysis Console (TAC) 4.0 software. Sample signals and statistical measurements were exported from TAC 4.0 software. Genes measured by multiple probes were individually numbered. Clustering was performed in Cluster 3.0 using log transformed data and genes were clustered using the uncentered correlation similarity metric and average linkage settings^59^. Heatmaps were generated using JavaTreeview^60^.

### Survival Analysis

Survival curves were generated using KM Plotter for Breast Cancer^34^ using probe 209434_s_AT for *PPAT* and 213892_S_AT for *APRT*. Patients were separated by upper and lower tercile of expression using the trichotomization option. Redundant samples were removed and biased arrays were excluded as per the default quality control settings.

### Cell lines and culture conditions

EMT and papillary tumor derived cell lines were cultured in Dulbecco’s Modified Eagle Medium (DMEM Corning, Corning, New York 10-017-CM) with 25 mM glucose without sodium pyruvate supplemented with 2 mM glutamine (Corning, 25-005-CI) 10% heat-inactivated fetal bovine serum (MilliporeSigma, Burlington Massachusetts, 12306C), and 1% penicillin and streptomycin (Corning, 30-002-CI). Cells were maintained at 37°C with 5% CO_2_.

### CRISPR/Cas9

Lentivirus mediated CRISPR/Cas9 genome editing was used to achieve gene knockout. Guide RNAs targeting *APRT* or *PPAT* were designed using the CRISPR-DO web application^62^. Plasmids containing dual guide RNA, puromycin resistance, and Cas9 co-expression were acquired from VectorBuilder. Plasmids containing scramble guide RNA, puromycin resistance, and Cas9 co-expression were also acquired from VectorBuilder. *APRT* KO dual guide RNA sequences are guide A) 5’-GTCGATCTTGCCGCTGTGCG-3’ and guide B) 5’-GTGTGCTCATCCGGAAACAG-3’. *PPAT* KO dual guide RNA sequences are guide A) 5’-CATACGAGGTACGCCACCAC-3’ and guide B) 5’-TACGCGGTGCGAGATCCATA-3’ The non-targeting puromycin-resistant scramble guide RNA sequence is 5’-GCACTACCAGAGCTAACTCA-3’. Lentiviral envelope and packaging plasmids were acquired from addgene. The VSVG plasmid was a gift from Bob Weinberg (Addgene plasmid # 8454; http://n2t.net/addgene:8454; RRID:Addgene 8454). The psPAX2 plasmid was a gift from Didier Trono (Addgene plasmid # 12260; http://n2t.net/addgene:12260; RRID:Addgene 12260). To produce lentivirus, HEK293T cells seeded in 10-cm plates were transfected using lipofectamine 3000 (ThermoFisher Scientific, L3000015) with 10.0 μg lentivirus plasmids, 0.5 μg VSVG, and 5.0 μg psPAX2 plasmids. The following morning, fresh DMEM with 15% FBS and 1% P/S was added, and cells were grown for another 48 h to generate virus. For transduction with lentivirus, the recipient EMT and papillary cells were seeded in 10-cm plates and the supernatant of transfected HEK293T was collected and passed through 0.45 um PVDF syringe filter. 5 ml of the viral supernatant and 5 ml of fresh media were added to recipient cell plates with polybrene (Fisher Scientific, TR1003G) at a final concentration of 4 μg/ml. The cells were cultured for 24 h followed by addition of fresh DMEM medium supplemented with 10% FBS and treatment for 10 days with 2 μg/ml puromycin for selection. After transduction, cell culture media was supplemented with 50 uM nucleosides (adenosine, cytidine, guanosine, inosine, thymidine, and uridine) in DMSO across all conditions to provide extracellular nucleotides for cells with deficient *de novo* biosynthesis. The puromycin selected cells were then resuspended to a concentration of 5 cells/ml and seeded 1 cell/well on 96-well plates. Surviving clones were expanded and analyzed for successful gene knockout. Genomic DNA was extracted using DNeasy Blood and Tissue Kit (Qiagen) to check for successful gene editing. The following primer pairs were used for PCR expansion and sequencing (marked with *) of *APRT* guide A: 5’-GGGTCACTCTCCTGTCCTTG-3’ and 5’-AGGACAGAGCAGAGTTCGTC-3’*, *APRT* guide B: 5’-GAGCTGTTCAGAAGGCAGGT-3’* and 5’-AGCGTTTCTGGGTGGTGTAA-3’, *PPAT* guide A: 5’-CTCAGGACGGTCAAGGCTAC-3’* and 5’-AAGATGCCTTTTGTCGGAGA-3’, and *PPAT* guide B: 5’-GCATACACCCCTCCTCAAGA-3’* and 5’-CATCAGAGACTGGCATAAGACG-3’. Tracking of Indels by Decomposition (TIDE) was used to evaluate successful gene editing^35^.

### Western blot analysis

Cell lysis and Western blot analysis were carried out according to standard protocols. The following dilutions of primary commercial antibodies were used as probes: 1:250 dilution of anti-APRT (Thermo Scientific, PA576741), 1:500 dilution of anti-PPAT (Proteintech 15401-1-AP), 1:1000 dilution of anti-vinculin (Cell Signaling Technology, E1E9V). Anti-APRT and anti-vinculin antibodies were diluted in 5% bovine serum albumin and incubated overnight at 4 °C. The anti-PPAT antibody was diluted in 5% milk and incubated for 60 minutes at room temperature per manufacturer recommendations. Secondary anti-rabbit antibodies (Cell Signaling Technology, 7074S) were diluted in 5% non-fat milk at a dilution of 1:1000 and incubated at room temperature for 1 h. Blots were imaged by chemiluminescence after incubation with Clarity Western ECL substrate (Bio-Rad, 1705061) using a ChemiDoc Imaging system (Bio-Rad).

### Isotope labeling studies

For isotope labeling experiments, DMEM without glucose or glutamine was prepared from powder (MilliporeSigma, D5030) and supplemented with ^13^C6-glucose (Cambridge Isotope Laboratories, Tewksbury, Massachusetts, CLM-1396) and unlabeled glutamine (MilliporeSigma, G8540). Labeled media was prepared with 10% dialyzed FBS (Sigma-Aldrich, F0392). Cells were then seeded and cultured as described above. Fresh cell culture media without nucleoside supplementation was added to cells for 1 hour prior to switching to isotope containing media. Prior to metabolite extraction, media was switched to isotope containing media and samples were collected at T = 240 minutes. Metabolite extraction and analysis were performed as above. Labeling data was corrected for natural isotope abundance using IsoCor^63^.

### In vivo tumor studies

To generate tumors, monoclonal KO cell lines were injected in 50 µl of a 1:1 mixture of DMEM:Matrigel (Corning, 354262) at 500,000 cells/50 µl into the fourth mammary fat pad of syngeneic 6 to 8 week old FVB mice. The resulting tumors grew to a size of 15 mm as measured by external calipers along the longest axis, at which time the tumors were harvested and fragmented into 3 mm pieces that were cryopreserved in a mixture of 90% FBS and 10% DMSO. Cryopreserved tumors were then thawed, washed in saline, and cut into 1-2mm fragments for implantation into the fourth mammary fat pad of recipient mice. These re-implanted tumors were then measured by external caliper 3 times weekly starting at 7 days post implantation until the experimental endpoint at 24 days post implantation. Tumor size was calculated as cross-sectional area using measurements from the longest and shortest axes. Mice were monitored for humane endpoints throughout the experiment according to institutional guidelines. At 24 days the tumors were collected, and a cross section of each tumor was formalin fixed for histological preparation.

### Histological analyses

All histological preparation and immunohistochemical staining was performed by the Investigative HistoPathology Laboratory at Michigan State University. Ki67 staining was measured using multiple images taken from distinct, non-necrotic regions of each tumor and evaluated as follows. For each tumor, at least 4 color images from distinct regions were acquired using an Olympus BX41 microscope operated at 10x magnification and saved as TIFF image files. Image processing was performed in ImageJ 1.52p (Fiji distribution). The color images were first deconvoluted into H (hematoxylin) and DAB (diaminobenzidine) color channels using Color Deconvolution (‘H DAB’ deconvolution matrix). Deconvoluted H and DAB images were saved as new TIFF images. For each image, smoothing was applied 5 times, then Auto Local Threshold was performed using Bernsen’s algorithm (window size 15, contrast threshold 15) to detect stained nuclei. Stained nuclei were counted using Analyze Particles (minimum size 150, minimum circularity 0.3). The above steps were looped over all images. To check that threshold parameters were appropriate, several output images were manually inspected to confirm that visually identifiable nuclei were properly counted. The percent Ki67 + nuclei was calculated as the ratio of DAB-stained nuclei counts (representing proliferating cells) to H-stained nuclei counts (representing all cells) for each image, and averaged across all images for each experimental group. TUNEL assays were evaluated using a single image of the full tumor cross section to determine the proportion of necrotic area to non-necrotic area of each tumor. Images were acquired using a Leica M165FC stereo microscope operated at 1x magnification and saved as TIFF image files. TUNEL assay images were also processed using ImageJ. Images were duplicated and color thresholding was used to select either the TUNEL + area (image 1) or the entire tumor area (image 2). The percent TUNEL + area was calculated as the ratio of image 1 area to image 2 area for each tumor and averaged across all tumors within each experimental group.

### Statistical analyses

Statistical analyses were performed using unpaired Student’s t-test except where otherwise noted. *p* values were adjusted in R using the p.adjust() function to account for multiple hypothesis testing using the Benjamini-Hochberg procedure (metabolites) or Hommel procedure (tumor measurements). All error bars presented are standard deviation. All figures except survival curves and heatmaps were generated using GraphPad Prism.

## Acknowledgments

The authors thank Deanna Broadwater, Elliot Ensink, and Hyllana Medeiros for helpful discussions and critical reading of this manuscript. The authors thank Eran Andrechek for providing primary MMTV-Myc EMT and MMTV-Myc papillary tumors. The authors also thank the MSU Mass Spectrometry and Metabolomics Core and the MSU Investigative HistoPathology Laboratory. **Funding:** This work was supported by the Office of the Assistant Secretary of Defense for Health Affairs, through the Breast Cancer Research Program, under Award No. W81XWH-15-1-0453 to SYL. This work was also supported by the Spectrum Health MD/PhD Fellowship and the Aitch Foundation Graduate Fellowship to MPO.

## Author Contributions

MPO performed metabolic profiling, gene expression analysis, survival analysis, cell culture, CRISRP/Cas9, Western blots, isotope labeling, and in vivo tumor studies. STT performed IHC studies and assisted with interpretation of results. SYL conceived, designed, and supervised the study. All authors contributed to writing the manuscript and have critically read, edited, and approved the manuscript.

