## Supplementary Material for "Targeting subtype-specific metabolic preferences in nucleotide biosynthesis inhibits tumor growth in a breast cancer model"

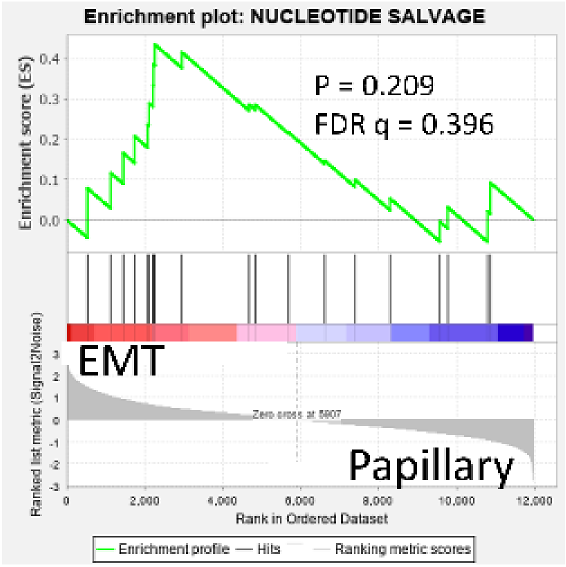


**Supplementary Figure 1: Gene set enrichment analysis for nucleotide salvage genes.** GSEA for nucleotide salvage genes are not significantly enriched in the EMT subtype.

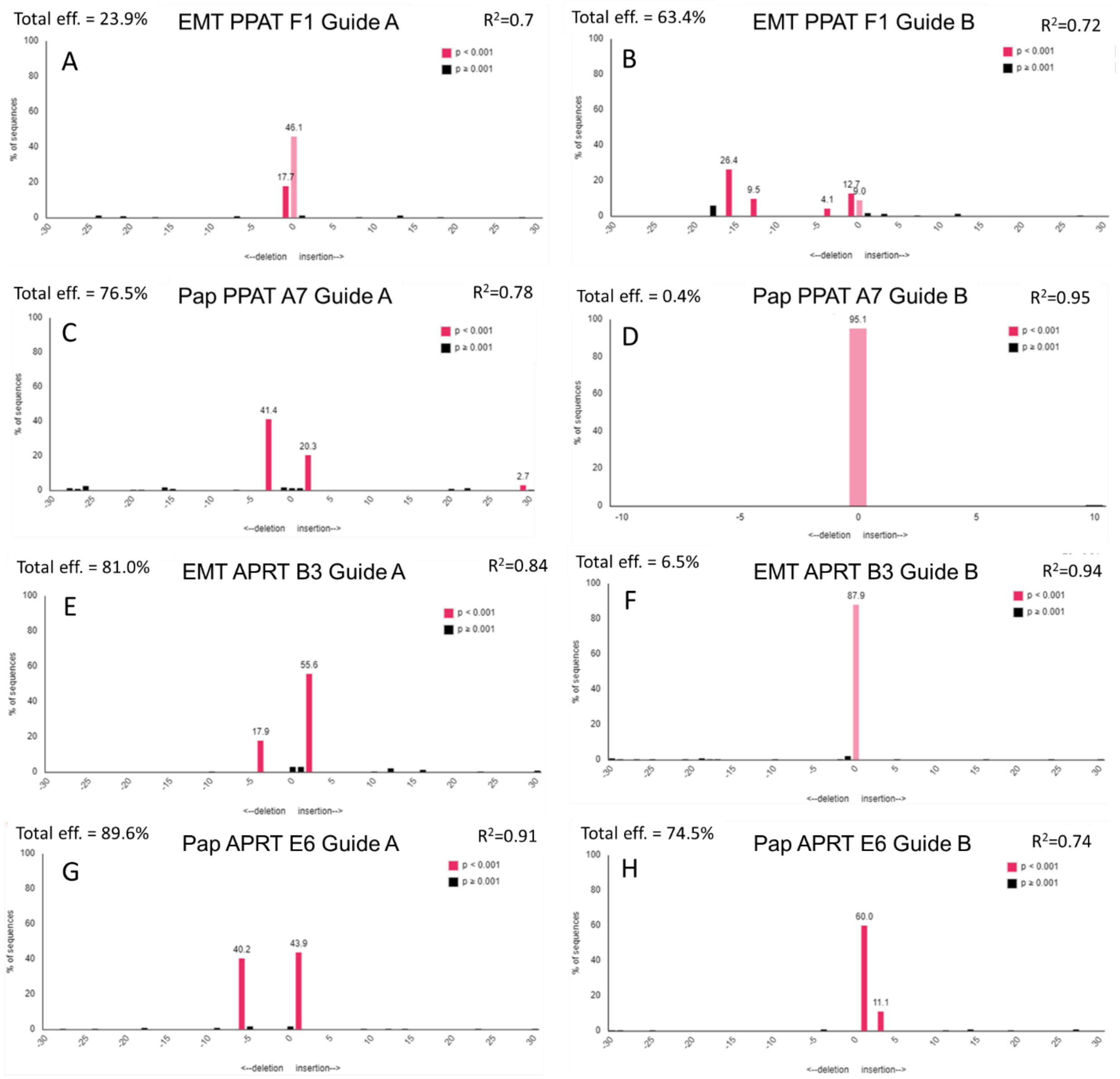


**Supplementary Figure 2: Gene editing verification of PPAT and APRT editing**. Sequencing of PPAT KO cell lines were validated using TIDE analysis. Pink bars denote insertion/deletion events with high confidence (p <0.001) for EMT PPAT KO clone F1 in (A) guide A and (B) guide B and for papillary PPAT KO clone A7 in (C) guide A and (D) guide B. Sequencing was also used to validate EMT APRT KO B3 in (E) guide A and (F) guide B and for papillary APRT KO E6 in (G) guide A and (H) guide B.

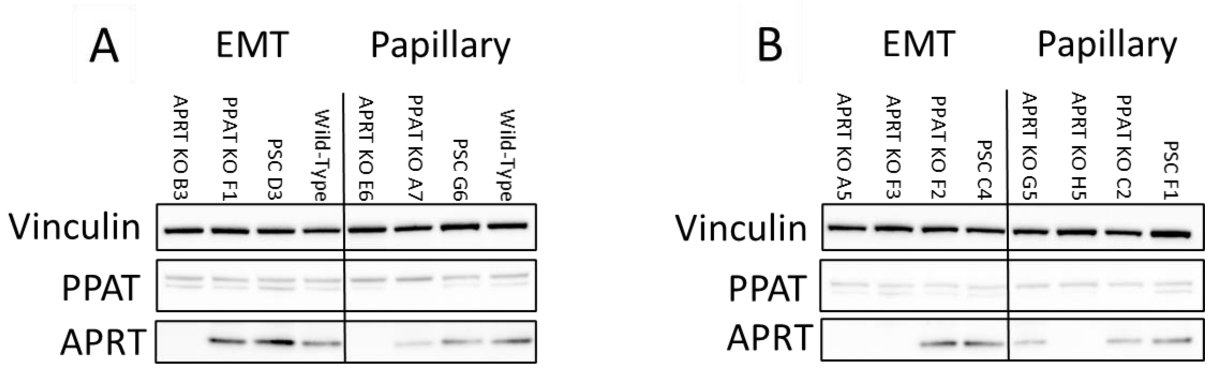


**Supplementary Figure 3: Protein verification of KO cell lines.** Western blotting was used to verify protein levels of (A) EMT APRT KO B3, PPAT KO F1, PSC D3, wild-type, papillary APRT KO E6, PPAT KO A7, PSC G6, and wild-type. Protein levels of (B) additional cell lines EMT APRT KO A5, APRT KO F3, PPAT KO F2, PSC C4, papillary APRT G5, APRT KO H5, PPAT KO C2, and PSC F1 were also verified.

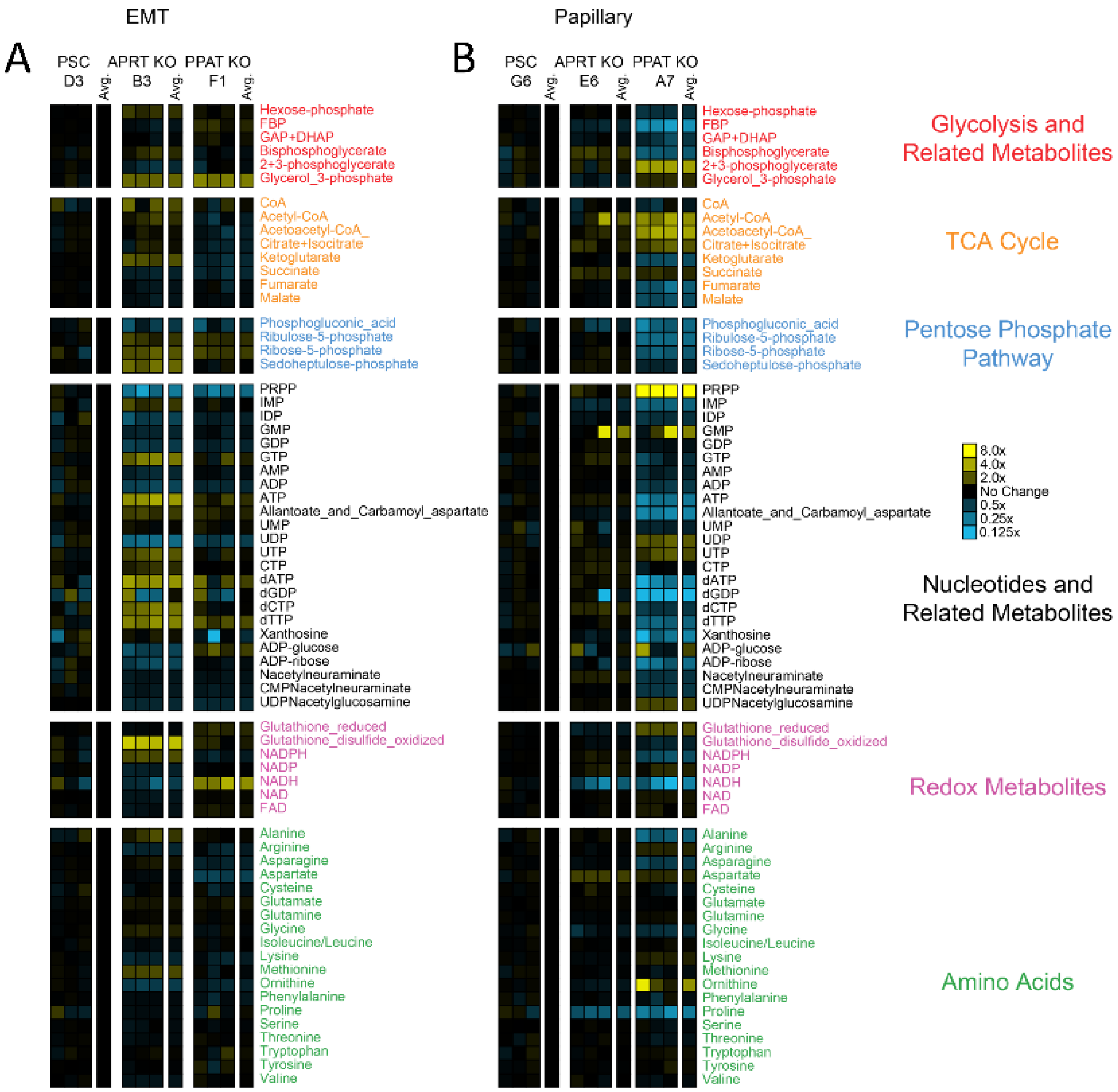


**Supplementary Figure 4: Full metabolic profiles of control and KO cell lines.** Heatmap indicating relative metabolite differences between control and KO cell lines in the (A) EMT and (B) papillary subtypes. Boxes indicate metabolite levels relative to the average of the PSC control for each subtype. Statistical comparisons are listed in **Supplementary Table 4**.

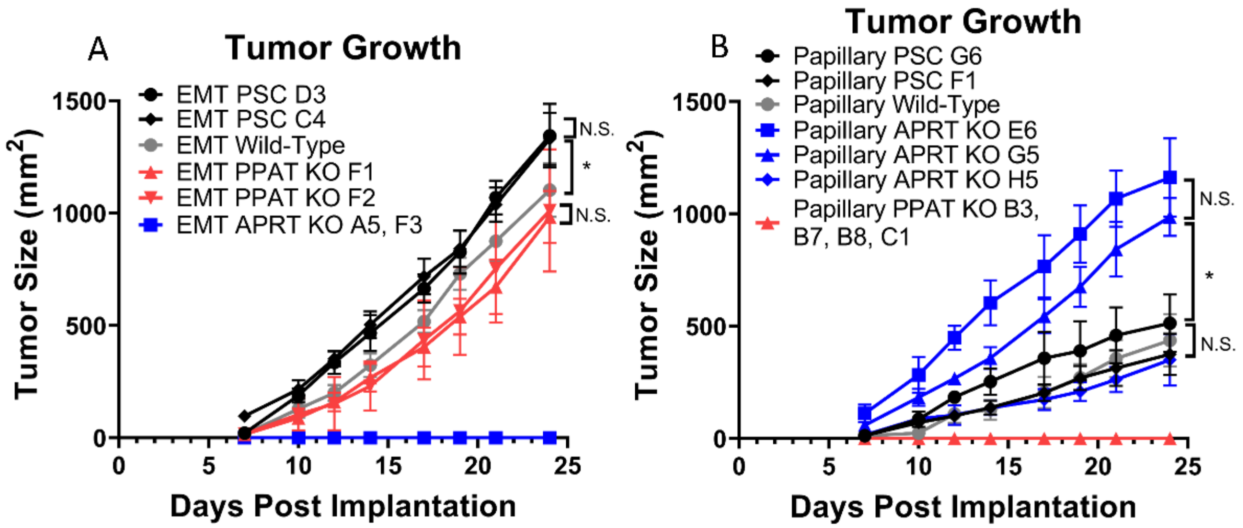


**Supplementary Figure 5: Tumor growth of additional clones.** In vivo growth curves for (A) EMT and (B) papillary tumors. Data are displayed as means ± S.D. (*p-value < 0.05). Statistical comparisons are listed in **Supplementary Table 5**.

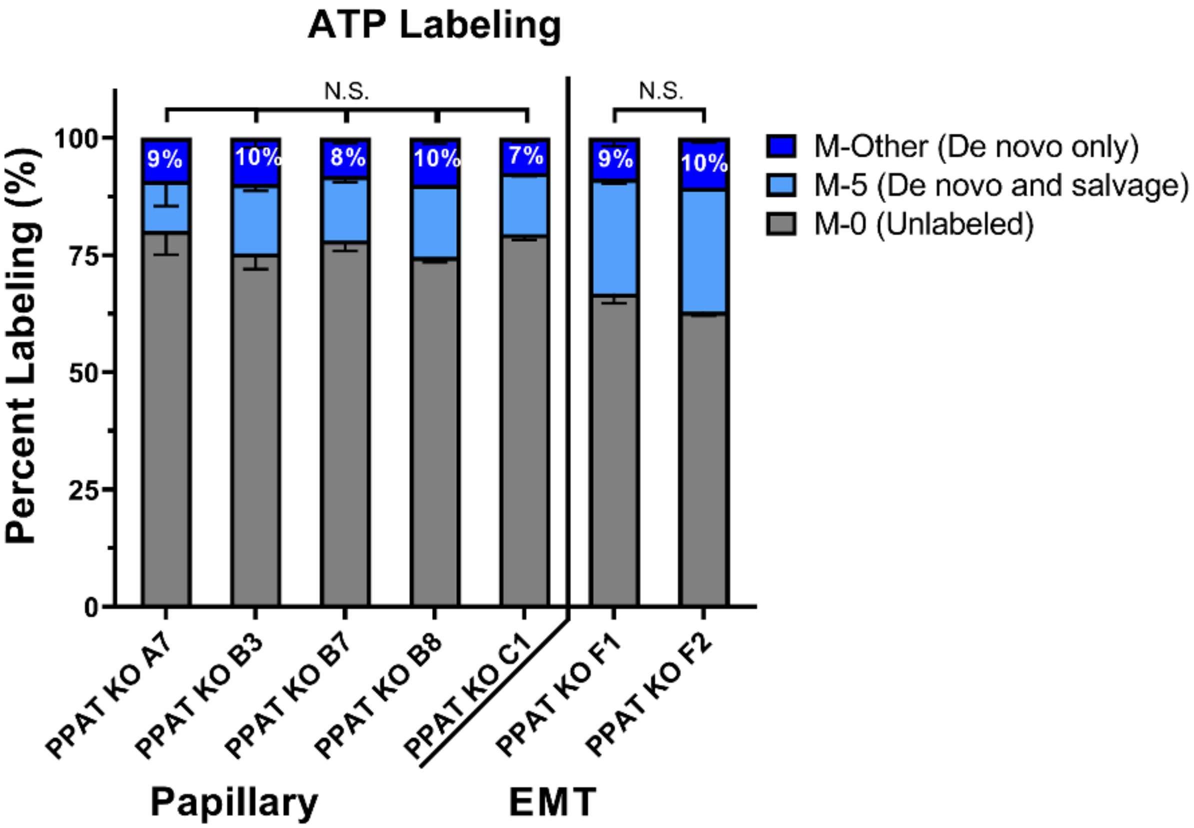


**Supplementary Figure 6: $^{13}$C-Isotope incorporation from glucose into ATP biosynthesis in additional PPAT KO clones.** Grey boxes represent the unlabeled (M-0 isotopologue) proportion of ATP. Light blue boxes represent the M-5 isotopologue, which can be derived from either de novo or salvage pathways. Dark blue boxes represent the sum of all other isotopologues of ATP (M1-4 and M6-10), which are derived from de novo ATP biosynthesis. Statistical comparisons are listed in **Supplementary Table 8**.

### SUPPLEMENTARY TABLE TITLES

**Supplementary Table 1. Metabolite abundance with statistical significance for Figure 1B.**

**Supplementary Table 2. Gene expression with statistical significance for Figure 2A.**

**Supplementary Table 3. $^{13}$C-Isotope percent labeling from glucose with statistical significance for Figure 4.**

**Supplementary Table 4. Metabolite abundance with statistical significance for Figure 5 and Supplementary Figure 4.**

**Supplementary Table 5. Tumor size with statistical significance for Figure 6 and Supplementary Figure 5.**

**Supplementary Table 6. Ki67 statistical significance for Figure 7 A-B.**

**Supplementary Table 7. TUNEL assay statistical significance for Figure 7 C-D.**

**Supplementary Table 8. $^{13}$C-Isotope percent labeling from glucose with statistical significance for Supplementary Figure 6.**
